# Mechanistic Interpretability of Fine-Tuned Protein Language Models for Nanobody Thermostability Prediction

**DOI:** 10.64898/2025.12.22.695878

**Authors:** Taihei Murakami, Yuki Hashidate, Yasuhiro Matsunaga

## Abstract

**Motivation:** While Protein Language Models (PLMs) fine-tuned on biophysical data achieve high predictive accuracy, the physical principles underlying their predictions remain obscure. Deciphering these representations offers a unique opportunity to not only interpret model decisions but also to discover novel biophysical insights governing protein properties. Here, we present a framework using Sparse Autoencoders (SAEs) to extract mechanistic knowledge from PLMs fine-tuned for nanobody thermo-stability.

**Results:** We fine-tuned the ESM-2 model on the nanobody thermostability dataset, achieving superior performance compared to significantly larger state-of-the-art models. SAE analysis successfully decomposed the model’s dense embeddings into sparse, interpretable features without loss of predictive accuracy. We characterized these features through both global and local analyses. Global analysis revealed that stabilizing features predominantly activate on the molecular surface, optimizing charge distributions, whereas destabilizing features map to the hydrophobic core, reflecting packing defects. Local analysis identified specific interactions, including known factors like the VHH-tetrad and critical disulfide bonds, as well as novel stabilizing candidates. Free Energy Perturbation calculations confirmed that these novel candidates exhibit structural compatibility, avoiding the severe destabilization seen in neighboring residues. By revealing these multi-scale biophysical rules, our approach demonstrates that interpreting fine-tuned PLMs provides a physics-grounded guide for rational protein engineering.

**Availability:** The data and source code of the proposed method are available at GitHub (https://github.com/matsunagalab/paper_nanobody-thermostability-sae) and Zenodo (DOI: 10.5281/zenodo.18012027).

**Contact:** Yasuhiro Matsunaga

**Supplementary information:** Supplementary data are available at *Bioinformatics* online.

## 1 Introduction

Protein Language Models (PLMs) are deep learning models that treat amino acid sequences as natural language, learning sequence grammar and statistical patterns through large-scale self-supervised training. PLMs generate rich embeddings effective for downstream tasks such as structure prediction and functional annotation (Rives et al. 2021; Elnaggar et al. 2022; Lin et al. 2023; Su et al. 2024; Hayes et al. 2025). However, the internal representations captured by PLMs remain largely elusive, representing a critical gap in our understanding. Recent studies suggest that PLMs strongly reflect information such as evolutionarily conserved sequence motifs and residue-residue covariation/contacts (Zhang et al. 2024). Although PLMs pre-trained on diverse protein sequence datasets like UniRef (Suzek et al. 2015) retain broad knowledge, they may not fully capture the physicochemical properties such as thermostability.

Interpretability techniques for understanding the prediction mechanisms of deep learning models are particularly important in scientific applications, but simple methods such as attention visualization (Rao et al. 2020; Vig et al. 2021; Leem et al. 2022) cannot sufficiently capture individual features or their contributions to predictive performance. Recently, research group at Anthropic has introduced Sparse Autoencoders (SAEs) (Trenton et al. 2023) into the field of large language models as a powerful approach that reconstructs internal dense representations as more interpretable monosemantic features and disentangles “superposition” at the neuron level. Recently, this method was applied to PLMs (Simon and Zou 2025; Adams et al. 2025; Gujral et al. 2025), where SAEs were independently applied SAEs to pre-trained models, demonstrating that biologically meaningful features can be extracted. However, these studies mainly focused on general features encoded in pre-trained models, and task-specific concepts acquired through fine-tuning to predict specific properties have not yet been sufficiently explored.

Supervised Fine-Tuning (SFT) adapts pre-trained models to specific tasks using labeled data, emphasizing task-specific features to enhance performance (Chu, Narang and Siegel 2024; Lafita et al. 2024; Schmirler, Heinzinger and Rost 2024; Wang et al. 2025). However, this performance gain comes with a trade-off: the model’s decision-making process becomes even more opaque, potentially obscuring which features contribute to predictions. Mechanistically elucidating which statistical patterns PLMs learn from data and how these contribute to predicting physicochemical properties is crucial for evaluating performance, detecting biases, and potentially deepening our understanding of biomolecular physics and chemistry.

Among various biophysical tasks for proteins, thermostability is a critical property directly relevant to functional maintenance. For instance, nanobodies (VHHs) are valued for their small size and high antigen recognition (Muyldermans 2013; Alexander and Leong 2024), but their applications require thermostable designs. Conventional methods for evaluating this important property rely on either low-throughput experiments like differential scanning calorimetry (Akiba et al. 2019; Ikeuchi et al. 2021) or computationally expensive molecular dynamics (MD) simulations (Bekker, Ma and Kamiya 2019). Therefore, establishing machine learning methods that can rapidly and accurately predict melting temperature (Tm) from sequence alone is strongly needed (Blaabjerg et al. 2023; Li et al. 2023). Previous studies have shown that general thermostability prediction does not improve simply through increased pre-training or model size expansion of PLMs, suggesting that fine-tuning of PLMs with labeled data is essential for the thermostability prediction of nanobodies (Lin et al. 2023).

This study aims to mechanistically elucidate the features PLMs acquire when fine-tuned for nanobody thermostability. We fine-tuned a PLM for the nanobody thermostability prediction task and constructed a high-accuracy fine-tuned PLM. We then analyzed its internal representations by applying SAEs to decompose the embedding representations into interpretable sparse features. We successfully extracted features significantly correlated with Tm from the fine-tuned model. Analysis of features correlating with high Tm values revealed that they capture not only known factors like VHH-tetrad residues and disulfide bonds but also novel stabilizing factors, while features correlating with low Tm values capture hydrophobic core residue groups critical for stability. This study represents that a combination of supervised fine-tuning and SAEs serves as an effective computational protocol for mechanistic interpretability. This approach bridges the gap between powerful predictive models and scientific insights by enabling the extraction of task-specific concepts from fine-tuned PLMs.

## 2 Methods

### 2.1 Datasets

For SFT, we utilized NbThermo (Valdés-Tresanco et al. 2023), a publicly available database, as the labeled dataset. NbThermo is a database that systematically collects nanobody Tm values and amino acid sequences manually curated from the literature, containing 567 nanobody entries.

For SAE training, we used sequence data extracted from INDI (Integrated Nanobody Database for Immunoinformatics) (Deszyński et al. 2022). INDI is a dataset integrating over 11 million nanobody sequences from various NGS data. We performed clustering at 70% sequence identity using Linclust (Steinegger and Söding 2018) to remove redundancy from this dataset, then randomly extracted 100,000 sequences for use as the SAE training dataset.

### 2.2 Supervised fine-tuning of Protein Language Model

In this study, we used the pre-trained ESM-2 8M model (consisting of 6 layers with embedding dimension *D*_*ESM−hidden*_ = 320) as the base model. We performed SFT on this model for the regression task of predicting Tm values from nanobody amino acid sequences (Fig. 1). The model architecture consists of ESM-2 with an added a linear regression head. The input to the linear head is a vector obtained by mean-pooling the embedding along the sequence length *N*_*aa*_ direction from the final layer (layer 6). Special tokens, <cls> and <eos>, were excluded from this calculation. First, as a baseline, we trained only the linear head using Ridge regression while keeping the ESM-2 weights frozen. The regularization parameter of Ridge regression was optimized through cross-validation. Hereafter, we refer to this baseline prediction model as the *pre-trained model*.

**Fig. 1.**
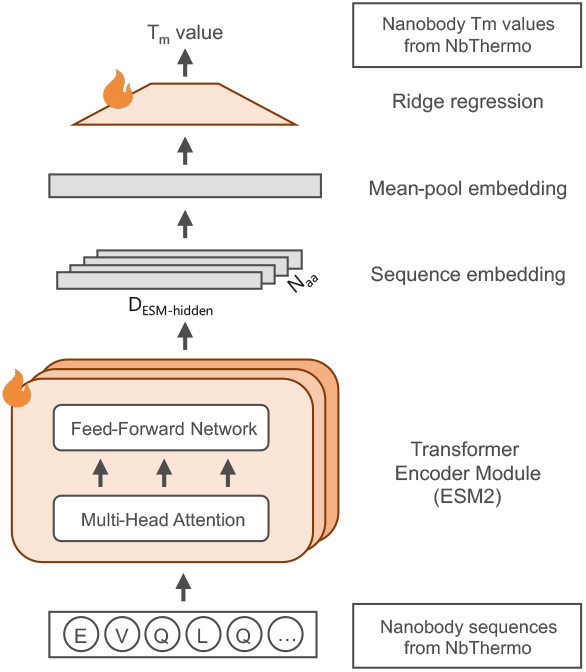
Schematic of the supervised fine-tuning of a protein language model for nanobody thermostability prediction. Architecture of the predictive model. Nanobody sequences are processed by the ESM-2 encoder. The mean-pooled embeddings are fed into a linear head to predict Tm. Both the encoder and linear head weights are updated using the NbThermo dataset.

Subsequently, we made both the ESM-2 main body and linear regression head weights updatable. We split the 567 entries in the NbThermo dataset into train:validation = 80:20 (train = 453, validation = 114). For the split, we used the same random seed as used in the TEMPRO model (Alvarez and Dean 2024) to enable quantitative comparison with it. For training, we used mean squared error (MSE) as the loss function and optimized the weights of both the ESM-2 encoder and prediction head for 50 epochs. We explored multiple conditions for hyperparameters such as learning rate, regularization coefficient, and batch size, adopting the settings that mostly minimized MSE. Hereafter, we refer to this SFT prediction model as the *SFT model*. Model performance evaluation was performed on the remaining 20% test data from the NbThermo dataset, using RMSE, MAE, R2 and Pearson correlation coefficient as metrics.

### 2.3 Training of Sparse Autoencoder

To interpret the internal representations, we applied SAEs with the same architecture as Simon and Zou (Fig. 2a). SAEs transform the dense residue embedding representation 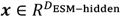 into a higher-dimensional *D*_SAE−hidden_ ≫ *D*_ESM−hidden_ but sparse feature vector 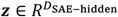, and reconstruct the original embedding representation from these sparse features. Specifically, the SAE encoder transforms ***x*** to the sparse latent vector ***z***, and the decoder reconstructs the embedding representation ***x***′ using dictionary vectors 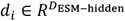 embedding (*i* = 1, … , *D*_SAE−hidden_) with the following equations:

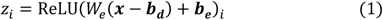

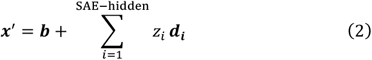

where *W*_*e*_ is a weight matrix, and ***b***_***d***_, ***b***_***e***_, ***b*** are bias terms. SAE training optimized reconstruction error (MSE) plus L1 norm of latent vector (sparse representation) ***z*** as a regularization term. In this study, we processed 100,000 sequences extracted from INDI with the fine-tuned model and used the embedding from the final layer (layer 6) as training targets for the SAE. The SAE hidden layer dimension was set to *D*_SAE−hidden_ = 10240 (32x expansion of *D*_ESM−hidden_ = 320), and training was performed with an L1 norm coefficient of 0.05 after hyperparameter tuning. As a result, the sparsity of ***z*** was approximately 2%.

**Fig. 2.**
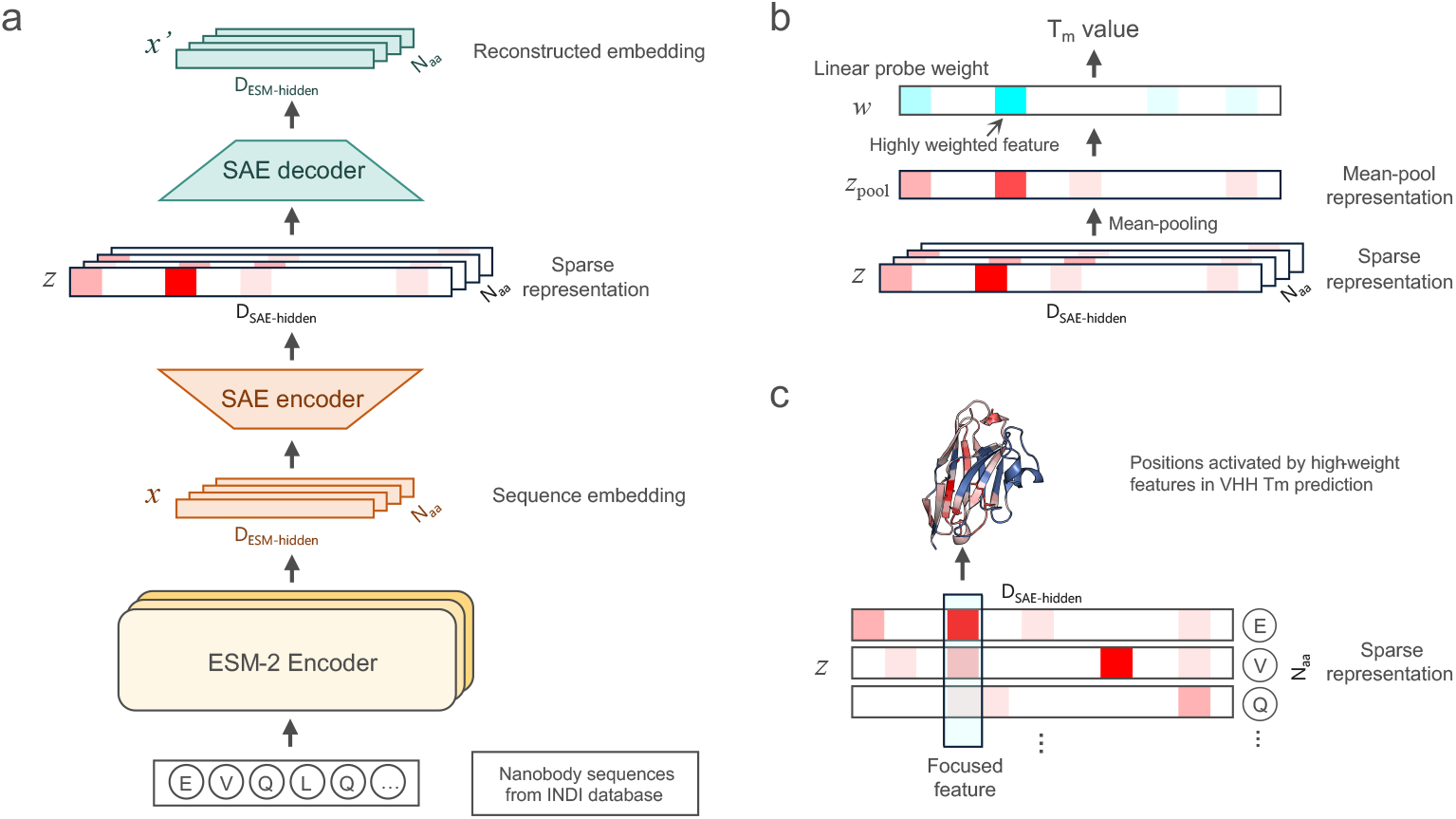
Methodology for interpreting the embedding representation of the fine-tuned model using a Sparse Autoencoder (SAE). **a**. Training of the SAE. The embedding representation ***x***, extracted from the final layer of the ESM-2 encoder for unlabeled nanobody sequences, are used to train an SAE. The SAE learns to reconstruct the dense embedding representation ***x***′ from a high-dimensional, sparse representation ***z***, effectively decomposing the original embedding into a linear combination of interpretable features. **b**. Identifying thermostability-related features. A linear model is trained to predict Tm values directly from the mean-pooled sparse representations ***z***_pool_. The learned weights *w* of the linear probe indicate the importance of each feature for the prediction task; features with large positive or negative weights are considered highly relevant to thermostability. **c**. Visualization and interpretation of features. For a feature identified as important, the specific amino acid residues that cause its activation are identified. These activated residues are then mapped onto the 3D structure of the nanobody to analyze their biophysical and structural significance.

To identify features with large contributions to thermostability, we added a linear head to the latent vector ***z*** (Fig. 2b) (Adams et al. 2025). In this approach, since the prediction head is linear, features with large absolute weight values in the linear head can be interpreted as important for Tm value prediction. In the calculation of prediction, we first input each sequence from the NbThermo dataset to the trained SAE to obtain sparse feature representations. We then mean-pooled the obtained feature representations ***z*** along the sequence length *N*_*aa*_ direction, aggregating each sequence into a single vector ***z***_pool_ of *D*_SAE−hidden_ = 10,240 dimensions. The linear head was trained using this vector as input via Ridge regression.

To verify whether the SAE truly obtained sparse representations of the underlying pre-trained model, we compared predictions of the SFT model with those of the pre-trained model. Both models’ predictive calculations used the NbThermo test dataset. Subsequently, for NbThermo sequences, particularly focusing on sequences with high and low Tm values, we investigated the biophysical significance by visualizing amino acid residues firing in features with large absolute weights in the SAE’s linear head. For structural visualization, we performed structure prediction from sequences using AlphaFold 3 (Abramson et al. 2024).

For interpreting the SAE sparse representation in terms of structure, we visualized how the SAE-extracted features contribute to nanobody thermostability with position-by-position detail along AHo numbering across the entire dataset. All 567 NbThermo sequences were aligned using AHo numbering, with gaps treated as zero. For each feature *k* with |*w*_*k*_| ≥ 0.05, we computed the per-position linear contribution as *w*_*k*_ × *z*_*k*_(*p*), where *w*_*k*_ denotes the Ridge weight learned from mean-pooled SAE features and *z*_*k*_(*p*) is the raw activation of feature *k* at Aho-numbering position *p*. We then averaged the contributions across the selected features separately for positive (*w*_*k*_ > 0, stabilizing) and negative (*w*_*k*_ < 0, destabilizing) sets to obtain the position-wise profiles. For structural visualization, we mapped these profiles onto AlphaFold 3 structure models of the highest-Tm sequence (for the positive map) and the lowest-Tm sequence (for the negative map) in the dataset.

### 2.4 Free energy perturbation

To quantify the thermodynamic contributions of specific residues identified by the SAE, we performed alchemical free energy perturbation (FEP) calculations. All molecular dynamics simulations for FEP were carried out using NAMD (Phillips et al. 2020). We utilized a customized version of the AlaScan plugin (Ramadoss et al. 2016) for VMD (Humphrey et al. 1996) to support mutations to arbitrary amino acids other than standard alanine scanning. The relative free energy changes (ΔΔG) were calculated using the Bennett Acceptance Ratio (BAR) method (Bennett 1976) implemented in the ParseFEP plugin (Liu et al. 2012) to ensure statistical convergence and accuracy.

## 3 Results

### 3.1 SFT improves prediction and acquires representations reflecting nanobody thermostability

To evaluate the effect of SFT of PLMs on nanobody thermostability prediction, we compared the pre-trained model (model with only the linear head trained) against the SFT model (model where all weights including EMS-2 portion fine-tuned for labeled datasets). While the pre-trained model achieved a Pearson correlation coefficient of 0.787, R2 of 0.609, RMSE = 6.621 °C, and MAE = 4.947 °C between predicted and measured Tm values, the SFT model achieved better Pearson correlation coefficient = 0.865, R^2^ = 0.739, RMSE = 5.402 °C, and MAE = 3.734 °C, demonstrating higher prediction accuracy (Fig. 3a). These results support that, as shown in previous studies (Schmirler, Heinzinger and Rost 2024), fine-tuning the entire model including the PLM improves task-specific prediction performance compared to training only the regression head.

**Fig. 3.**
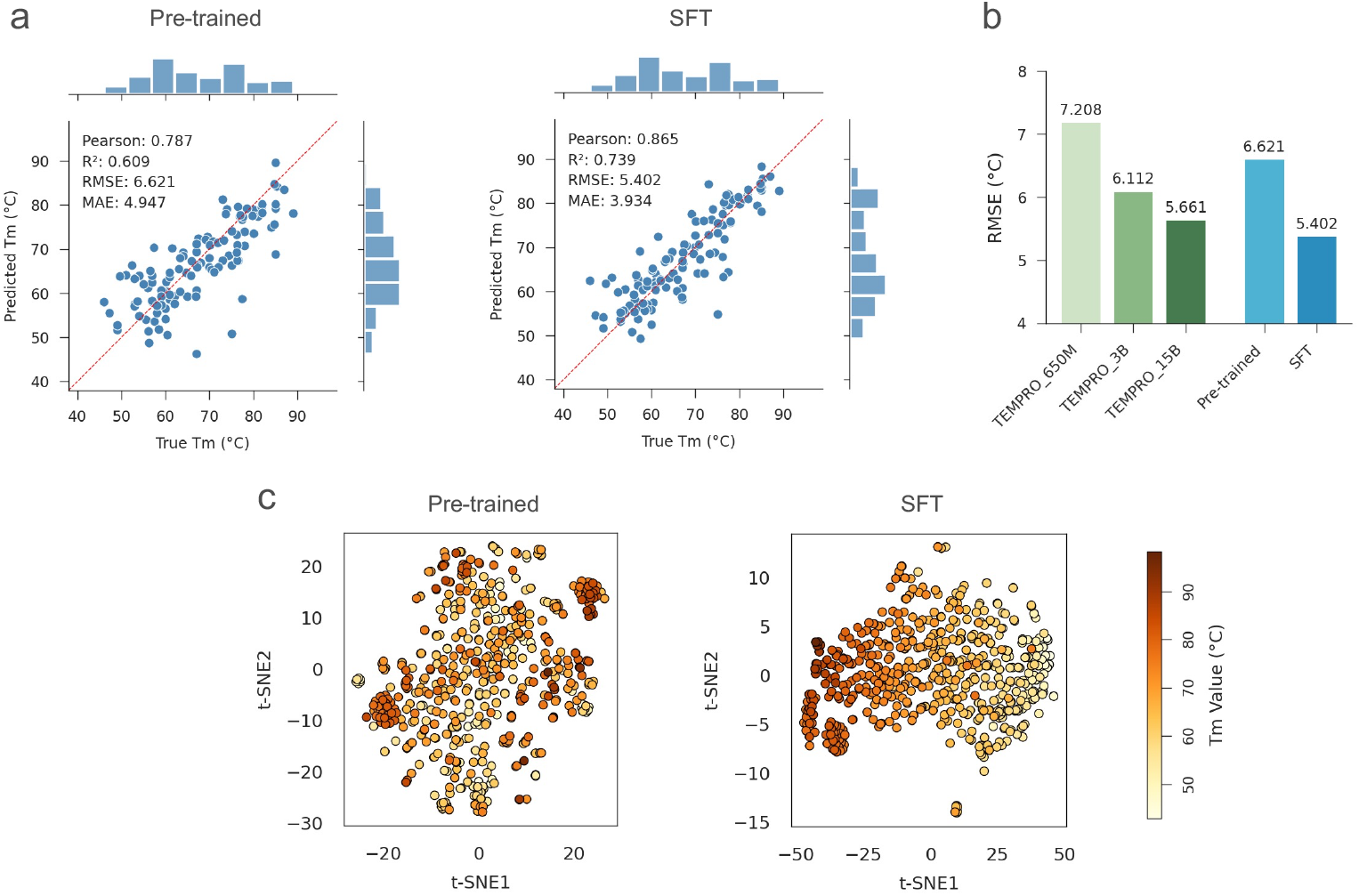
Supervised fine-tuning (SFT) improves thermostability prediction. **a**. Comparison of prediction performance on the test set between the pre-trained model (left), where only the final regression head is trained, and the SFT model (right), where the entire model is fine-tuned. Scatter plots show predicted Tm versus true Tm. **b**. Benchmarking of model performance. The Root Mean Square Error (RMSE) of our pre-trained and SFT models (based on ESM-2 8M) is compared with the reported performance of various sizes of the state-of-the-art TEMPRO model (based on ESM-2 650M, 3B, and 15B). **c**. t-SNE visualization of the embeddings representation from the final layer of the pre-trained (left) and SFT (right) ESM-2 encoders. Each point represents a nanobody sequence from the test and training sets, colored by its experimental Tm value.

Furthermore, we compared the performance of our constructed model with TEMPRO, one of the existing state-of-the-art models for nanobody thermostability prediction. TEMPRO consists of an architecture with a prediction head composed of multilayer perceptrons (MLP) attached to the output of the ESM-2 encoder. The ESM-2 encoder weights are frozen and not fine-tuned with Tm data. TEMPRO shows gradual improvement in Tm value prediction performance by expanding model size, achieving RMSE = 7.208 °C, 6.112 °C, and 5.661 °C for models based on ESM-2 650M, 3B, and 15B models, respectively. While TEMPRO’s largest model (TEMPRO_15B using ESM-2 15B) achieves RMSE = 5.661 °C, our SFT model (based on ESM-2 8M) demonstrated superior RMSE = 5.402 °C despite its smaller model size (Figure 3b). This result indicates that the PLM with SFT is more effective for improving downstream task prediction performance than simply expanding model scale.

Next, to investigate whether the SFT model extracts sequence patterns related to thermostability, we performed t-SNE visualization of the embedding representations from the final layer (layer 6) of the ESM-2 encoder (Fig. 3c). While no structure correlated with Tm values was observed in the embedding space of the pre-trained model, the SFT model showed embedding representations clearly correlated with Tm values. Specifically, in the SFT model, a strong correlation (Pearson’s r = - 0.93) was observed between the horizontal axis (t-SNE1) in the space and Tm values, indicating that thermostability-related features are captured as representations.

### 3.2 SAE of the SFT model disentangles important features for nanobody thermostability

Through our analysis thus far, we found that the SFT model has acquired effective embedding representations for predicting thermostability. Next, to unravel what specific features comprise these internal representations, we used SAEs to disentangle the embedding representations from the final layer (layer 6) of the ESM-2 encoder with sparse representations. We applied SAEs to both the pre-trained and SFT models for comparison to verify whether the SFT model’s SAE truly extracts features important for thermostability prediction.

First, we evaluated whether the sparse representations obtained by SAE could sufficiently approximate the ESM-2 embedding representations. We added a linear regression model predicting Tm values using sparse representations as input and examined whether its prediction performance was equivalent to the original linear regression model using the embedding representations as input (Fig. 4a for the pre-trained model, and Fig. 4c for the SFT model). In the SFT model, the prediction accuracy from sparse representations was RMSE = 5.574 °C, quantitatively similar to the prediction accuracy from the original dense embedding representations (RMSE = 5.402 °C). Furthermore, the correlation of predicted values was extremely high at Pearson = 0.998 (Fig. 4c), indicating that the sparse representations almost completely approximate the information related to Tm prediction. Similarly, in the pre-trained model, the Tm prediction accuracy from sparse representations showed RMSE = 5.906 °C and high correlation with predictions from the original embeddings (Pearson correlation = 0.95) (Fig. 4a).

**Fig. 4.**
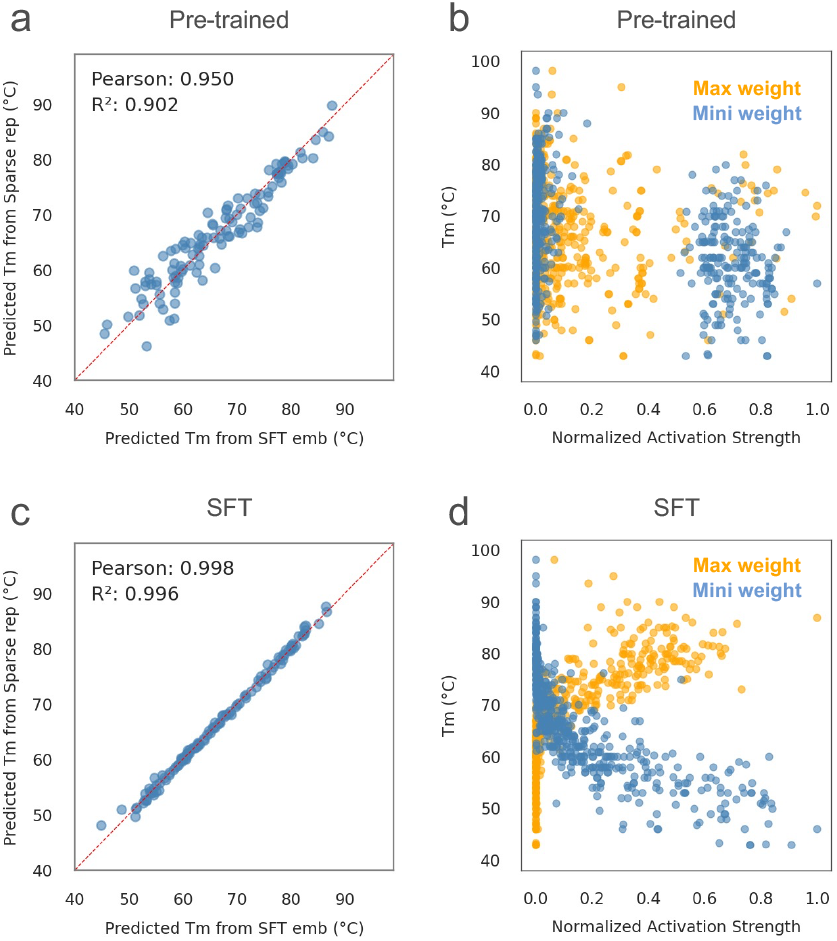
Analysis of the pre-trained model. **a**. The Tm values predicted from the sparse representation are highly correlated with those predicted from the original dense embedding, showing the SAE preserves information. **b**. The activation strengths of the most important positive (maximum weight) and negative (minimum weight) features against the true Tm values. **c-d**. Analysis of the SFT model.

Next, to investigate whether the SFT model’s sparse representations truly succeed in disentangling features related to thermostability, we examined the relationship between feature activation strengths (in the sparse representation) and Tm values for components with large weights in the linear model (Figs. 4b and 4d). Specifically, we identified the maximum and minimum weight components in the linear model and examined the relationship between activation strength (the elements of ***z***_<==>_ in Fig.2) and Tm values. In the SFT model, the feature with the maximum positive weight showed good correlation between Tm and activation strength, while the feature with the most negative weight showed good negative correlation between Tm and activation strength (Fig. 4d and Fig. S1). This indicates that the sparse representations disentangled from the SFT model retain important features for nanobody thermostability and instability. In contrast, the pre-trained model showed no clear correlation like that seen in the SFT model (Fig. 4b and Fig.S2). This suggests that the pre-trained model’s ESM-2 encoder does not acquire biophysical concepts specialized for Tm, having embedding representations mixed with various protein properties learned through the masked language model task.

### 3.3 Global pattern interpretation of SAE features

We visualized how SAE features globally contribute to nanobody stability across all sequences. For the entire NbThermo sequences, we aggregated the activation value at each Aho-numbering position weighted by the linear probe model and obtained the averaged activations at each position over the sequences. This averaged profile was separately calculated for positive weights (stabilizing) and negative weights (destabilizing), respectively (Fig. 5a). We further mapped these activations onto AlphaFold 3 predicted structures of the sequences with the highest and lowest Tm in the dataset to examine their spatial distribution (Fig. 5b).

**Fig. 5.**
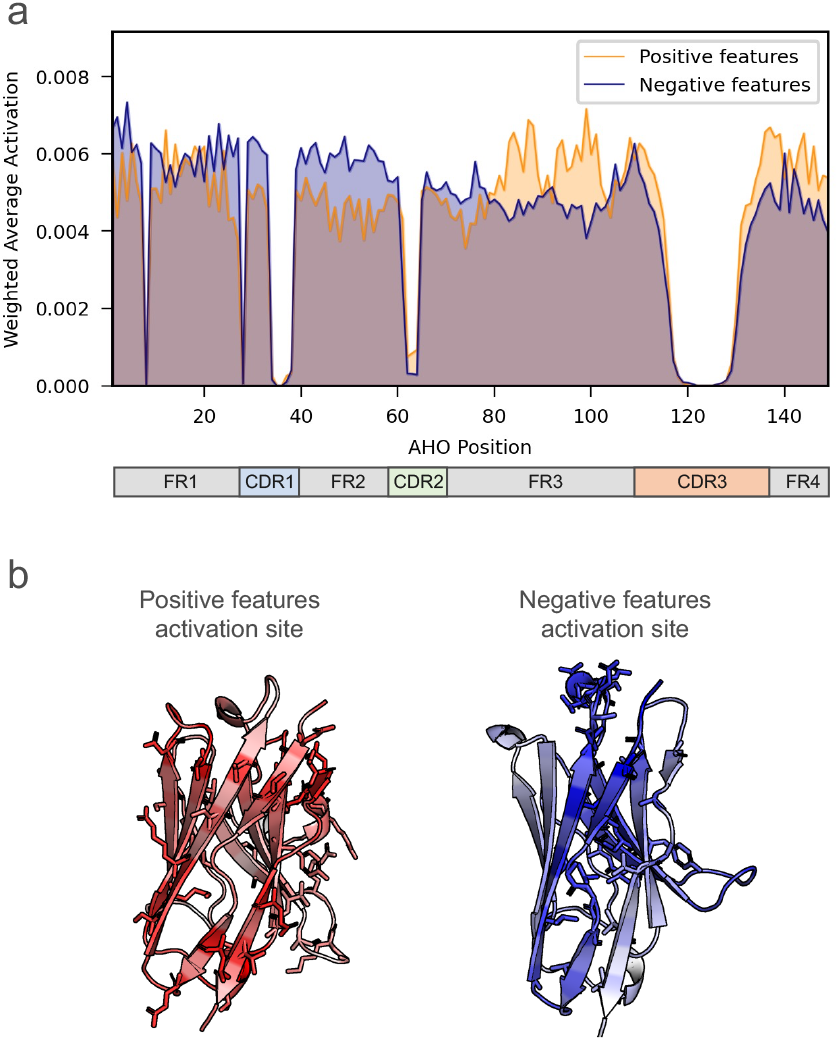
Global mapping of SAE-derived feature contributions to nanobody stability. **a**. AHo-aligned, position-dependent distributions of the averaged activation, shown separately for stabilizing (positive-weight) and destabilizing (negative-weight) feature sets. Positive contributions peak broadly across FR3-FR4, while negative contributions are relatively higher and more dispersed across FR1-FR2. **b**. Structural views of the same contributions mapped onto AlphaFold 3 structural models of the highest-Tm (for positive features) and lowest-Tm (negative features) sequences in the dataset, illustrating surface-oriented patterns for stabilizing contributions and core-oriented patterns for destabilizing ones.

From Fig. 5a, the weighted activation of positive features is relatively higher in FR3 and FR4, whereas that of negative features is relatively higher from FR1 to FR2. In the structural mapping in Figure 5b, positive features show stronger activation at residues located on molecular surfaces, while negative features show stronger activation at residues located in the molecular interior. These observations suggest that, positively weighted features are related to the surface properties, including organized charge/polarity patterns and salt-bridge/hydrogen-bond networks contributing to increased thermal stability. By contrast, in the interior, hydrophobic-core packing defects or increased desolvation cost are likely to decrease thermal stability.

### 3.4 Local pattern interpretation of SAE features

We then analyzed local amino acid patterns identified by the SAE and mapped them onto nanobody model structures (Fig. 6). First, we examined amino acid residues firing in features with large positive weights of the linear model. As already shown in Fig. 4d, residues firing in this feature contribute to improving Tm values.

**Fig. 6.**
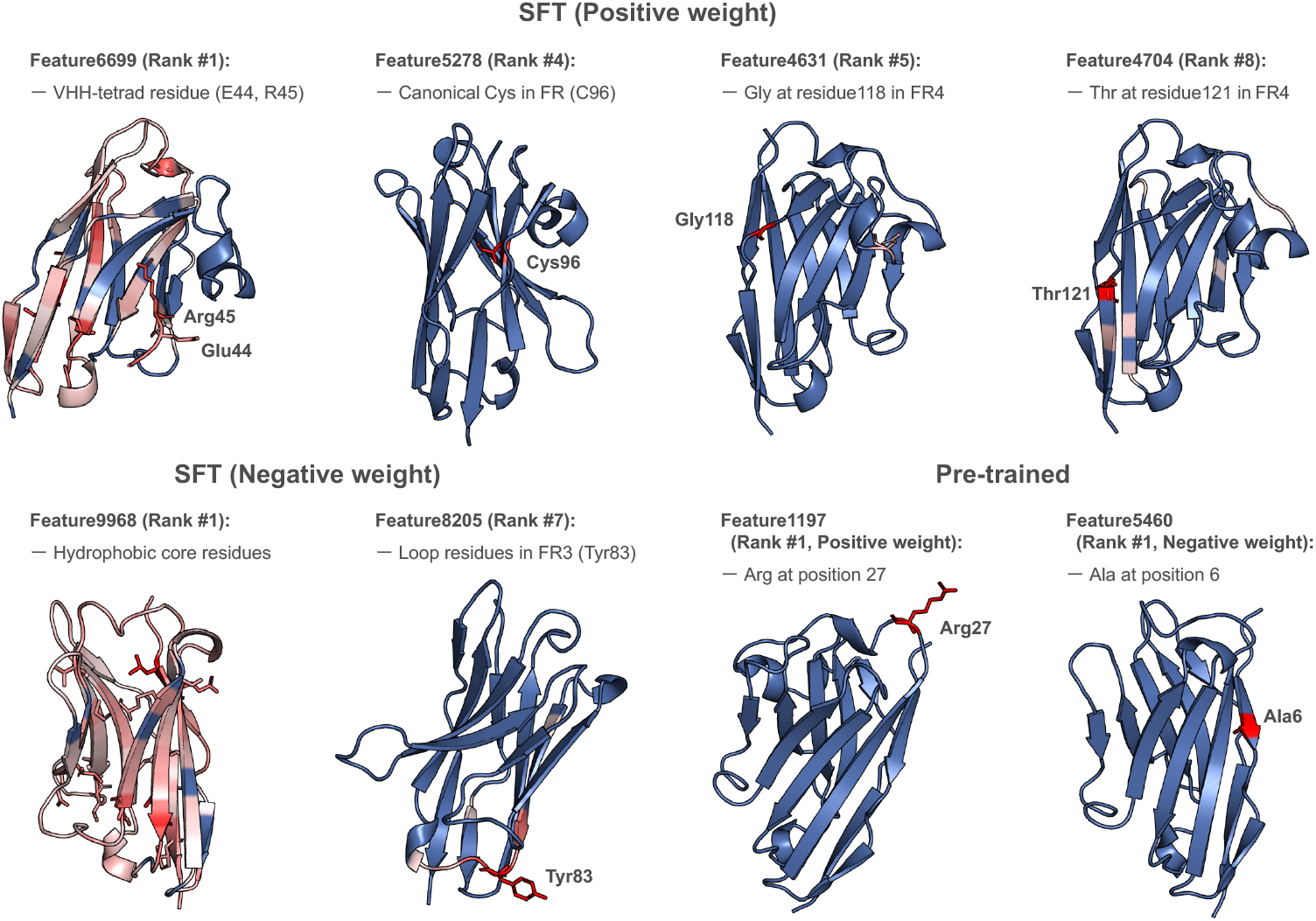
Structural interpretation of key features identified by the SAE in the SFT and pre-trained models. Residues that are highly activated by specific features are mapped in red onto a representative nanobody structure. (Top: SFT, Positive weight) Features that positively correlate with high Tm values (stabilizing features). These include Feature 6699 activating Glu44 and Arg45 (a key VHH-tetrad residue), Feature 5278 activating Cys96 (canonical framework disulfide bond). Feature 4631 activates Gly118, which permits the local twist required to connect FR4 β-sheets and thereby reduces conformational strain; and Feature 4704 activates Thr121, whose side chain points inward and can form a hydrogen bond with a nearby main chain. (Bottom: SFT, Negative weight) Features that negatively correlate with high Tm values (destabilizing features). Features 9968 activate residues distributed throughout the hydrophobic core, suggesting they represent the formation of this fundamental stability element. Feature 8205 activates FR3 loop residues (with Tyr83 most prominent), suggesting loop-induced strain that reduces stability. (Bottom: Pre-trained) For comparison, important features from the pre-trained model activate localized residues (Feature 1197: Arg27, Feature 5460: Ala6).

For the feature with the largest positive weight (Feature 6699), identifying residues in the highest-Tm sequence revealed Glu44 and Arg45 showed strong activation (here, the residue numbering is defined using the highest-Tm sequence). Glu44 and Arg45 are among the so-called VHH-tetrad positions (H37, H44, H45, H47), which are important substitutions where hydrophobic residues at the VH/VL interface in antibodies are replaced with hydrophilic residues in nanobodies (Muyldermans et al. 1994; Muyldermans 2021). This activation pattern was consistently observed in other high-Tm sequences as well (Fig. S5). These positions are known to be biophysically crucial residue groups for stabilizing nanobodies as a single-domain antibody (Rouet et al. 2015).

The feature corresponding to the fourth largest positive weight (Feature 5278) showed strong activation at Cys96. Cys96 forms a disulfide bond with Cys21 in framework region 1 (FR1), critical for nanobody stability (Liu, Schittny and Nash 2019). The features corresponding to the fifth and eighth largest positive weights (Feature 4631 and Feature 4704) showed particularly strong activation at Gly118 and Thr121, respectively. Although conserved in nanobodies, their contributions to thermostability remain elusive. We hypothesize that Gly118 contributes to stability by permitting the local twist required for connecting the β-sheets in FR4, thereby reducing global conformational strain; and that Thr121, which is oriented toward the protein interior, contributes to stability by forming a hydrogen bond between its side chain and a nearby main-chain group.

To test these hypotheses and validate the biophysical significance of the SAE-extracted features, we performed FEP calculations. We selected representative sequences from the NbThermo dataset that naturally lack these residues and possess relatively low Tm, making them suitable targets for testing stability improvements. Specifically, we used NbThermo ID 214 (modeled as seq217) and ID 426 (seq443) for Feature 4631; ID 340 (seq354) and ID 218 (seq221) for Feature 4704; and ID 335 (seq339) for Feature 8205.

For Feature 4631, the substitutions to Gly 118 (A106G in ID 214 and S113G in ID 426) yielded ΔΔG values of 1.81 kcal/mol and 0.96 kcal/mol, respectively. Similarly, for Feature 4704, substitutions to Thr 121 (I115T in ID 340 and V118T in ID 218) resulted in ΔΔG values of 0.27 kcal/mol and 0.40 kcal/mol. While these slightly positive values might suggest a lack of stabilization, they fall within a reasonable range considering the accuracy of state-of-the-art FEP calculations (Mean Absolute Error values ranging from 0.77 to 1.80 kcal/mol) (Koenekoop et al. 2025) and the tendency for inefficient sampling to bias towards instability. Indeed, in the vicinity of Gly118 (Feature 4631), neighboring mutations such as Y104G and T109G in ID 214 resulted in high energetic costs (5.32 kcal/mol and 3.32 kcal/mol, respectively), contrasting with the milder effect of the SAE-targeted mutation (1.81 kcal/mol). Similarly, for Thr121 (Feature 4704), neighboring mutations caused significant destabilization (e.g., G114T in ID 340: 6.63 kcal/mol; G117T in ID 218: 3.11 kcal/mol), whereas the SAE-targeted mutations remained neutral (0.27 kcal/mol and 0.40 kcal/mol). Overall, the FEP calculations suggest that the SAE does not merely identify simple “super-stabilizers,” but rather detects structurally optimized positions where specific residues fit seamlessly into the nanobody scaffold without the steric strain that plagues adjacent positions.

Next, we examined amino acid residues firing in features with large negative weights of the linear probe model. The seventh most negative feature (Feature 8205) exhibited localized activation in the FR3 loop, with Tyr83 showing the strongest activation. To investigate this, we performed FEP calculations on a sequence naturally containing Tyr83 (ID 335, modeled as seq339). The removal of Tyr (Y83S) resulted in a ΔΔG of 1.80 kcal/mol, indicating that Tyr83 itself contributes to local stability. However, this contribution is mild compared to the catastrophic destabilization observed upon mutating the adjacent residue L84S (6.82 kcal/mol). Another neighbor, N82S, showed a similar moderate value (1.63 kcal/mol). The SAE’s assignment of a negative weight to Tyr83 suggests that the model captures a broader context, perhaps identifying Tyr83 as a marker for lineages that are globally less thermostable or that require compensatory interactions, distinct from the universally critical hydrophobic core.

Apart from these, several of the remaining top-10 positive-weight features and top-10 negative-weight features lacked spatially localized activating residues and were difficult to interpret (Fig. S3)

In contrast to the SFT model, features important in the pre-trained model were qualitatively different from those captured by the SFT model, with consistently localized activation (Fig. 6 and Fig. S4). In the maximum positive weight feature (feature 1197), activation was concentrated at a single loop residue, Arg27; in the feature with the maximum negative weight (feature 5460), activation was concentrated at a single residue, Ala6. As shown in Fig. 4b, these pre-trained features do not correlate with nanobody thermostability.

Collectively, these findings highlight two key capabilities of local amino acid patterns identified by the fine-tuned PLM. First, by avoiding the severe destabilization observed in neighboring mutations, the model demonstrates it has learned structural permissibility, effectively identifying biophysically “safe zones” within dense protein packing. Second, by pinpointing compatible yet non-native residues, the SAE uncovers latent evolutionary optimization space. This suggests the model can identify stability-enhancing opportunities that nature may have bypassed due to complex evolutionary trade-offs, offering a guide for rational protein engineering.

## 4 Conclusions

In this study, we have demonstrated that a combination of SFT and SAE serves as an effective computational protocol for mechanistic interpretability for task-specific PLMs. Here by applying SAEs to the embedding of this SFT model, we decomposed them into interpretable sparse representations without losing predictive performance, capturing features related to thermostability. Feature analysis revealed that features correlating with high Tm values capture not only known factors like VHH-tetrad residues and disulfide bonds but also novel stabilizing factors, while features correlating with low Tm values capture hydrophobic core residue groups fundamental to stability. This approach provides a new pathway for mechanistically elucidating internal representations acquired by PLMs for specific biophysical properties, contributing to further performance improvements and rational protein design applications in the future.

While the fine-tuned PLMs identifies important residues, interpretation methods require further sophistication. Currently, identifying the biophysical meaning of individual features extracted by SAEs still heavily relies on manual experts analysis, as done in this study. To accelerate and more comprehensively perform this interpretation, developing interactive visualization systems like InterPLM (Simon and Zou 2025) would be effective. Future approaches could employ large language models to automatically annotate the function and structural roles from patterns of amino acid residues. However, ensuring reliability requires verification through energy calculations using MD simulations.

The insights gained from this study lead to promising future prospects. A key direction is extending the scope of this research beyond thermostability to other biochemical or biophysical data important in antibody-based drug development, such as aggregation propensity and binding affinity to other molecules. This would deepen understanding of universal or property-specific sequence motifs governing diverse protein properties

## Supplementary data

Supplementary data are available at *Bioinformatics* online.

## Author Contributions

Y.M., and T.M. wrote the manuscript. Y.M., Y.H, and T.M. developed the code for the method. Y.M., Y.H, and T.M. performed the analysis. All authors read and approved the final manuscript.

## Data and Code Availability

The data supporting the findings of this study have been deposited in the Zenodo repository (DOI: 10.5281/zenodo.18012027). The code is publicly available on GitHub (https://github.com/matsunagalab/paper_nanobody-thermo-stability-sae). Our implementation builds upon the sparse autoencoder (SAE) framework of Simon and Zou (Simon and Zou 2025) (see their original code at https://github.com/ElanaPearl/InterPLM) and incorporates only minor rather than major structural changes.

## Acknowledgements

We would like to thank Kentaro Sasaki (Saitama University) for helpful discussions on this work.

## References

1. Abramson J, Adler J, Dunger J et al. Accurate structure prediction of biomolecular interactions with AlphaFold 3. Nature 2024;630:493–500.

2. Adams E, Bai L, Lee M et al. From Mechanistic Interpretability to Mechanistic Biology: Training, Evaluating, and Interpreting Sparse Autoencoders on Protein Language Models. 2025, DOI: 10.1101/2025.02.06.636901.

3. Akiba H, Tamura H, Kiyoshi M et al. Structural and thermodynamic basis for the recognition of the substrate-binding cleft on hen egg lysozyme by a single-domain antibody. Sci Rep 2019;9:15481.

4. Alexander E, Leong KW. Discovery of nanobodies: a comprehensive review of their applications and potential over the past five years. J Nanobiotechnol 2024;22:661.

5. Alvarez JAE, Dean SN. TEMPRO: nanobody melting temperature estimation model using protein embeddings. Sci Rep 2024;14:19074.

6. Bekker G, Ma B, Kamiya N. Thermal stability of single – domain antibodies estimated by molecular dynamics simulations. Protein Science 2019;28:429–38.

7. Bennett CH. Efficient estimation of free energy differences from Monte Carlo data. Journal of Computational Physics 1976;22:245–68.

8. Blaabjerg LM, Kassem MM, Good LL et al. Rapid protein stability prediction using deep learning representations. eLife 2023;12:e82593.

9. Chu SKS, Narang K, Siegel JB. Protein stability prediction by fine-tuning a protein language model on a mega-scale dataset. PLOS Computational Biology 2024;20:e1012248.

10. Deszyński P, Młokosiewicz J, Volanakis A et al. INDI—integrated nanobody database for immunoinformatics. Nucleic Acids Research 2022;50:D1273–81.

11. Elnaggar A, Heinzinger M, Dallago C et al. ProtTrans: Toward Understanding the Language of Life Through Self-Supervised Learning. IEEE Trans Pattern Anal Mach Intell 2022;44:7112–27.

12. Gujral O, Bafna M, Alm E et al. Sparse autoencoders uncover biologically interpretable features in protein language model representations. Proc Natl Acad Sci USA 2025;122:e2506316122.

13. Hayes T, Rao R, Akin H et al. Simulating 500 million years of evolution with a language model. Science 2025;387:850–8.

14. Humphrey W, Dalke A, Schulten K. VMD: Visual molecular dynamics. Journal of Molecular Graphics 1996;14:33–8.

15. Ikeuchi E, Kuroda D, Nakakido M et al. Delicate balance among thermal stability, binding affinity, and conformational space explored by single-domain VHH antibodies. Sci Rep 2021;11:20624.

16. Koenekoop L, Van De Brug N, Jespers W et al. Accurate predictions of protein mutational effects accelerated with a hybrid-topology free energy protocol. Commun Chem 2025;8:362.

17. Lafita A, Gonzalez F, Hossam M et al. Fine-tuning Protein Language Models with Deep Mutational Scanning improves Variant Effect Prediction. 2024, DOI: 10.48550/arXiv.2405.06729.

18. Leem J, Mitchell LS, Farmery JHR et al. Deciphering the language of antibodies using self-supervised learning. Patterns 2022;3:100513.

19. Li M, Wang H, Yang Z et al. DeepTM: A deep learning algorithm for prediction of melting temperature of thermophilic proteins directly from sequences. Computational and Structural Biotechnology Journal 2023;21:5544–60.

20. Lin Z, Akin H, Rao R et al. Evolutionary-scale prediction of atomic-level protein structure with a language model. Science 2023;379:1123–30.

21. Liu H, Schittny V, Nash MA. Removal of a Conserved Disulfide Bond Does Not Compromise Mechanical Stability of a VHH Antibody Complex. Nano Lett 2019;19:5524–9.

22. Liu P, Dehez F, Cai W et al. A Toolkit for the Analysis of Free-Energy Perturbation Calculations. J Chem Theory Comput 2012;8:2606–16.

23. Muyldermans S. Nanobodies: Natural Single-Domain Antibodies. Annu Rev Biochem 2013;82:775–97.

24. Muyldermans S. A guide to: generation and design of nanobodies. The FEBS Journal 2021;288:2084–102.

25. Muyldermans S, Atarhouch T, Saldanha J et al. Sequence and structure of VH domain from naturally occurring camel heavy chain immunoglobulins lacking light chains. Protein Eng Des Sel 1994;7:1129–35.

26. Phillips JC, Hardy DJ, Maia JDC et al. Scalable molecular dynamics on CPU and GPU architectures with NAMD. The Journal of Chemical Physics 2020;153:044130.

27. Ramadoss V, Dehez F, Chipot C. AlaScan: A Graphical User Interface for Alanine Scanning Free-Energy Calculations. J Chem Inf Model 2016;56:1122–6.

28. Rao R, Meier J, Sercu T et al. Transformer protein language models are unsupervised structure learners. 2020:2020.12.15.422761.

29. Rives A, Meier J, Sercu T et al. Biological structure and function emerge from scaling unsupervised learning to 250 million protein sequences. Proc Natl Acad Sci USA 2021;118:e2016239118.

30. Rouet R, Dudgeon K, Christie M et al. Fully Human VH Single Domains That Rival the Stability and Cleft Recognition of Camelid Antibodies. Journal of Biological Chemistry 2015;290:11905–17.

31. Schmirler R, Heinzinger M, Rost B. Fine-tuning protein language models boosts predictions across diverse tasks. Nat Commun 2024;15:7407.

32. Simon E, Zou J. InterPLM: discovering interpretable features in protein language models via sparse autoencoders. Nat Methods 2025;22:2107–17.

33. Steinegger M, Söding J. Clustering huge protein sequence sets in linear time. Nat Commun 2018;9:2542.

34. Su J, Li Z, Han C et al. SaprotHub: Making Protein Modeling Accessible to All Biologists. 2024, DOI: 10.1101/2024.05.24.595648.

35. Suzek BE, Wang Y, Huang H et al. UniRef clusters: a comprehensive and scalable alternative for improving sequence similarity searches. Bioinformatics 2015;31:926–32.

36. Trenton Bricken, Adly Templeton, Joshua Batson, Brian Chen, Adam Jermyn, Tom Conerly, Nicholas L Turner, Cem Anil, Carson Denison, Amanda Askell, Robert Lasenby, Yifan Wu, Shauna Kravec, Nicholas Schiefer, Tim Maxwell, Nicholas Joseph, Alex Tamkin, Karina Nguyen, Brayden McLean, Josiah E Burke, Tristan Hume, Shan Carter, Tom Henighan, Chris Olah. Towards Monosemanticity: Decomposing Language Models With Dictionary Learning. https://transformer-circuits.pub/2023/monosemantic-features

37. Valdés-Tresanco MS, Valdés-Tresanco ME, Molina-Abad E et al. NbThermo: a new thermostability database for nanobodies. Database 2023;2023:baad021.

38. Vig J, Madani A, Varshney LR et al. BERTology Meets Biology: Interpreting Attention in Protein Language Models. 2021, DOI: 10.48550/arXiv.2006.15222.

39. Wang M, Patsenker J, Li H et al. Supervised fine-tuning of pre-trained antibody language models improves antigen specificity prediction. PLOS Computational Biology 2025;21:e1012153.

40. Zhang Z, Wayment-Steele HK, Brixi G et al. Protein language models learn evolutionary statistics of interacting sequence motifs. Proc Natl Acad Sci USA 2024;121:e2406285121.

