## supplementary data for "Mechanistic Interpretability of Fine-Tuned Protein Language Models for Nanobody Thermostability Prediction"

### S.1 Correlation analysis between feature activation and nanobody thermostability

We presented the relationship between activation strength and Tm values only for the features with the largest positive and negative weights (Fig. 4). To verify the generality of these observations, we extended this analysis to the top 10 features with the largest positive weights and the bottom 10 features with the largest negative weights for both the SFT and pre-trained models.

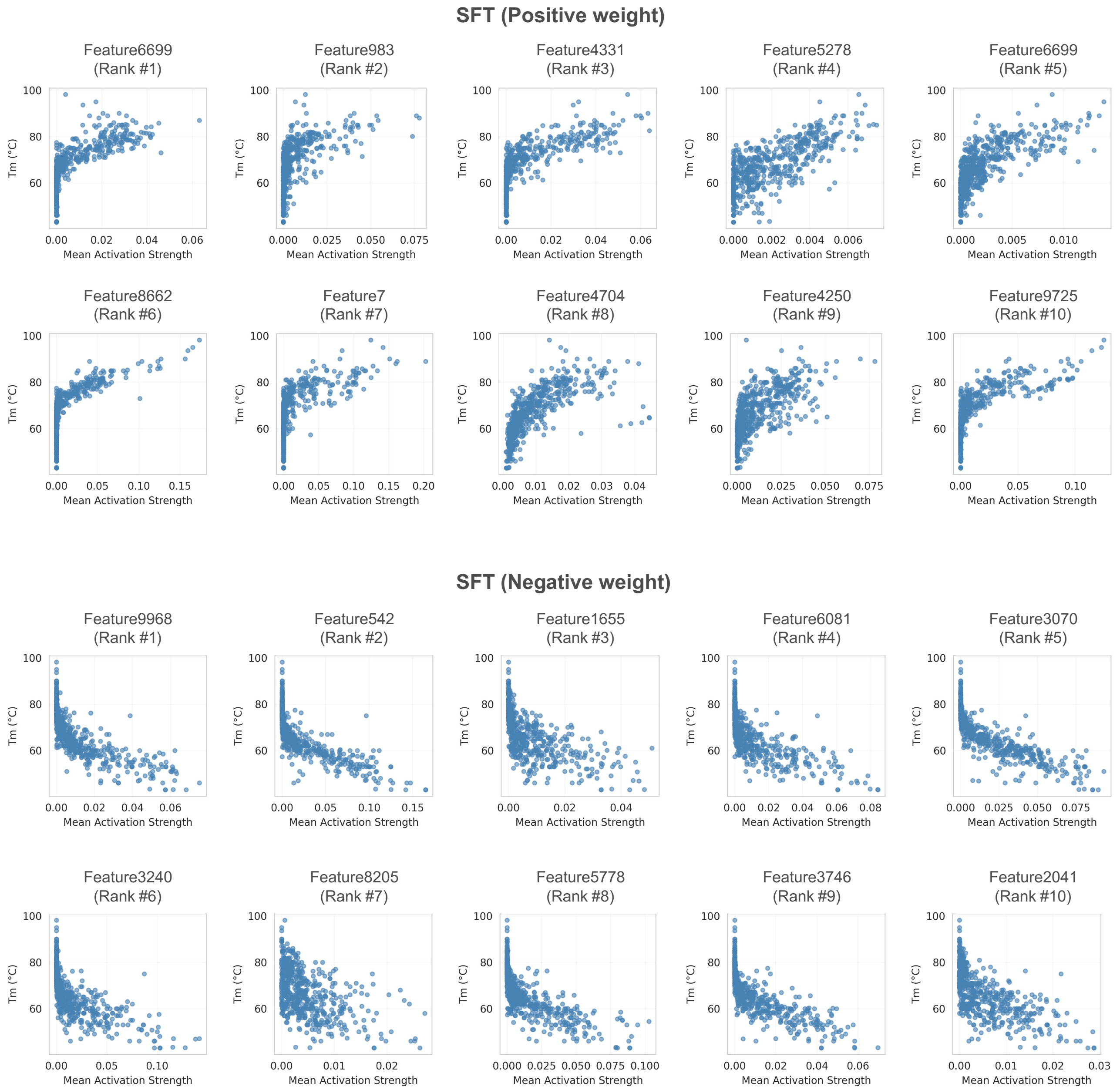


**Figure S1. Relationship between feature activation and thermal stability in the SFT model.** Scatter plots showing the normalized activation strength (x-axis) versus the true melting temperature (Tm) in the NbThermo dataset (y-axis). (Top: Positive weight) The 10 features with the largest positive weights (Rank #1-#10) in the linear regression head of the SFT model. (Bottom: Negative weight) The 10 features with the largest negative weights (Rank #1-#10) in the linear regression head of the SFT model.

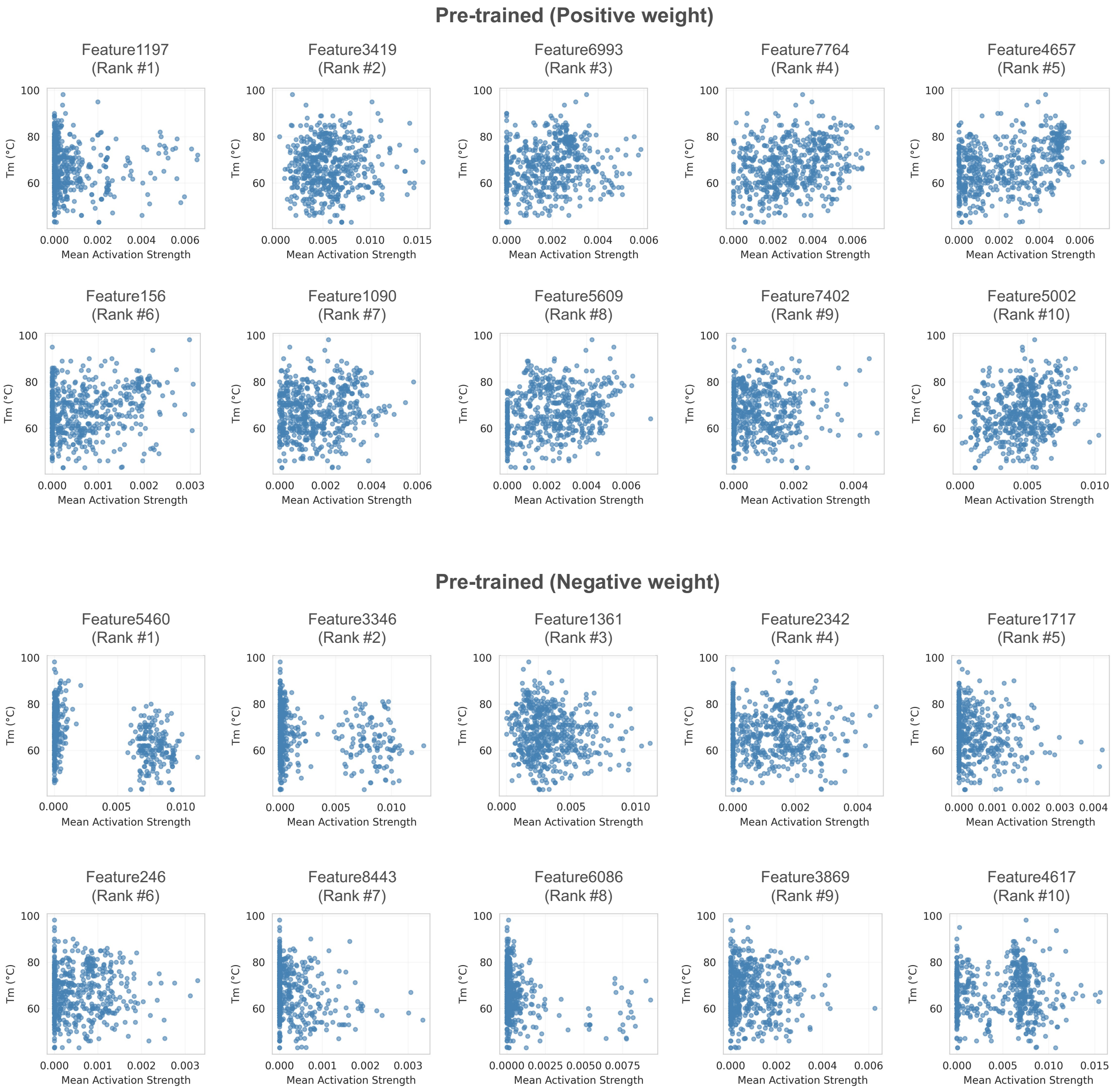


**Figure S2. Relationship between feature activation and thermal stability in the pre-trained model.** Scatter plots showing the normalized activation strength (x-axis) versus the true melting temperature (Tm) in the NbThermo dataset (y-axis). (Top: Positive weight) The 10 features with the largest positive weights (Rank #1-#10) in the linear regression head of the pre-trained model. (Bottom: Negative weight) The 10 features with the largest negative weights (Rank #1-#10) in the linear regression head of the pre-trained model.

### S.2 Structural mapping of the other features

For features with the largest positive or negative weights that were not shown in the main figures, we display their activation positions in nanobodies by highlighting high-activation residues and mapping them onto representative AlphaFold 3 models.

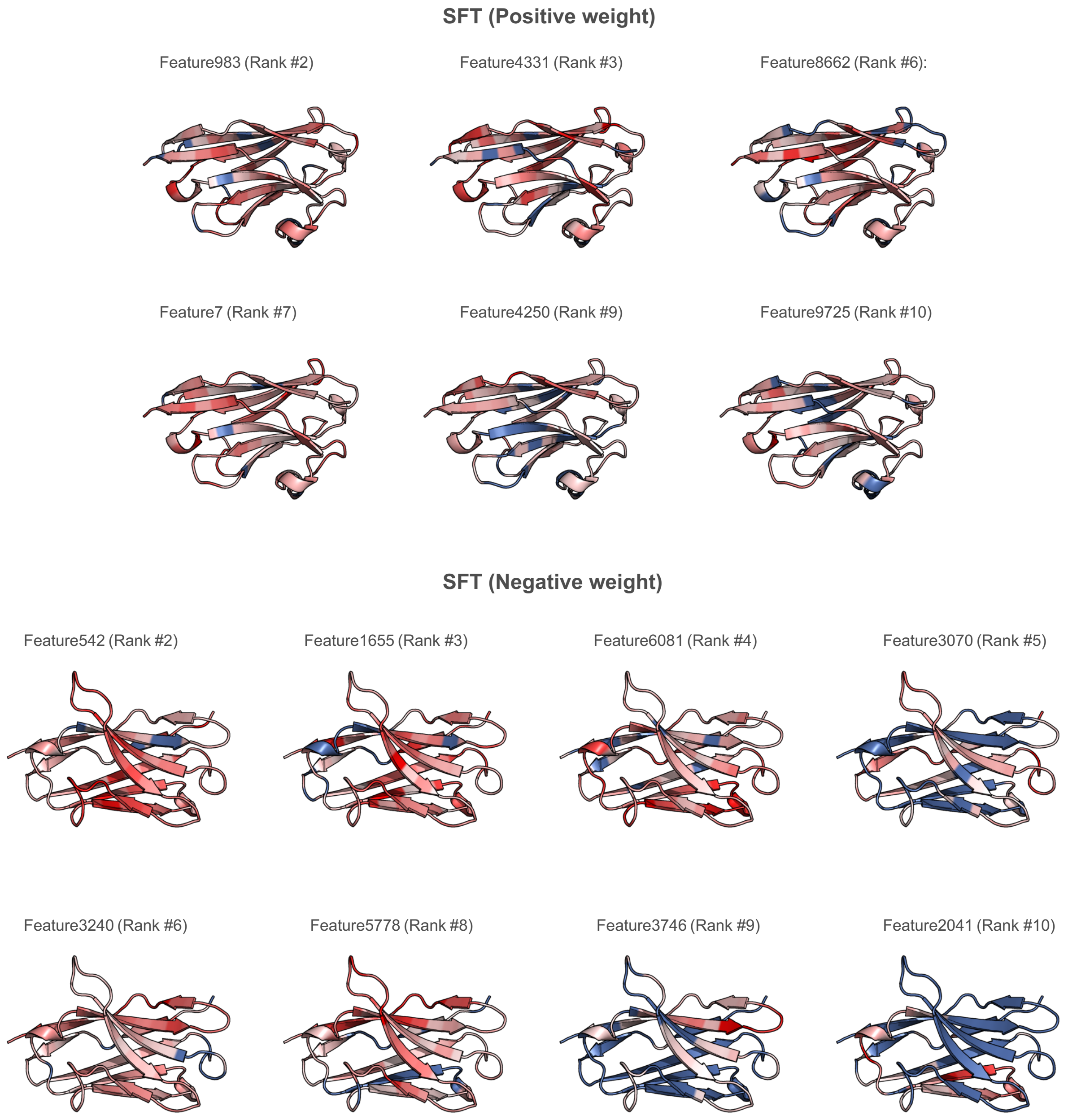


**Figure S3. Structural mapping positive features identified by the SAE in the SFT model.**

(Top: Positive weight) Positvely weighted features mapped on to the model structure of the highest-Tm sequence. (Bottom: Negative weight) Negatively weighted features mapped on to the model structure of the lowest-Tm sequence.

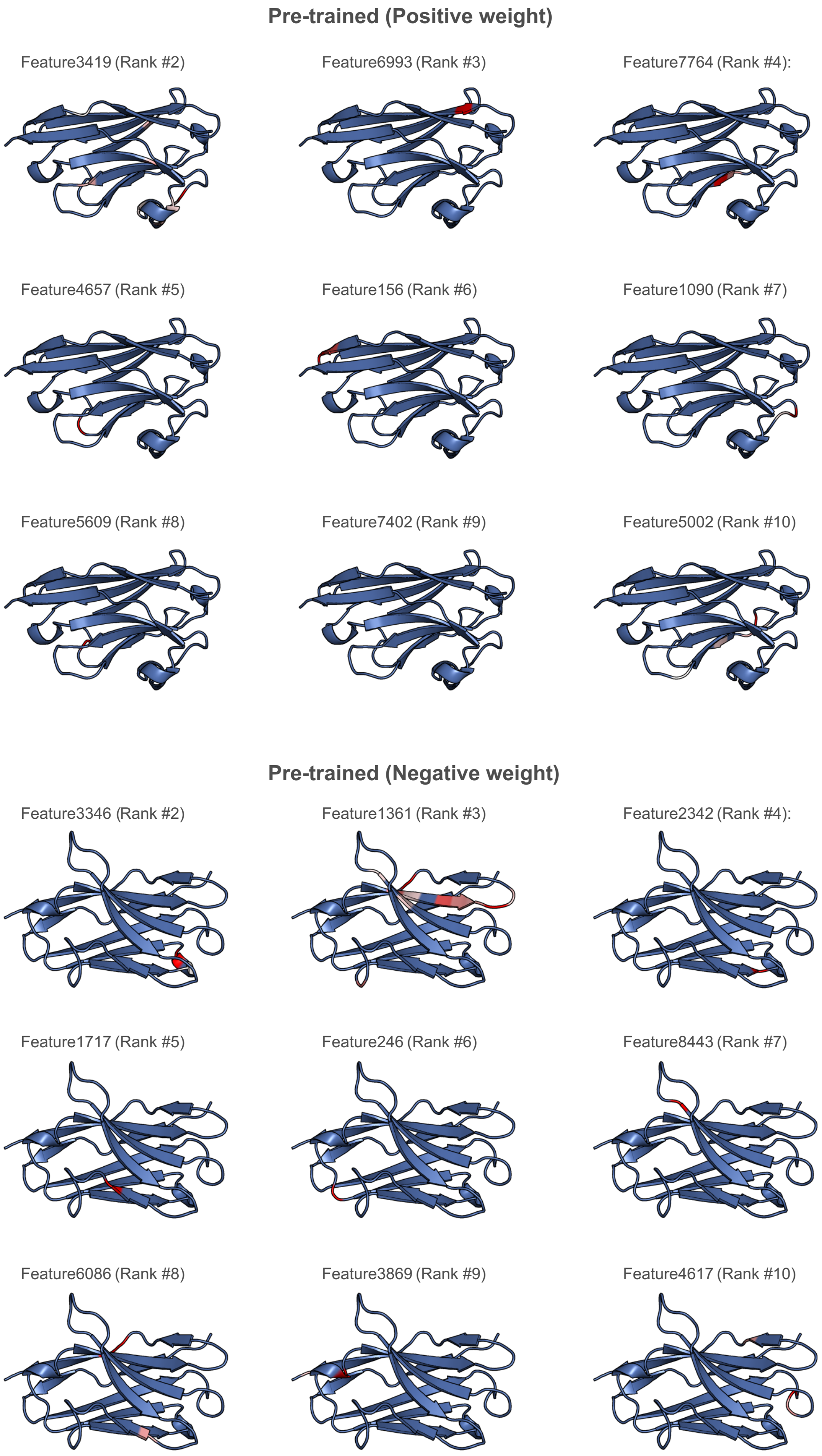


**Figure S4. Structural mapping negative features identified by the SAE in the pre-trained model.** (Top: Positive weight) Positvely weighted features mapped on to the model structure of the highest-Tm sequence. (Bottom: Negative weight) Negatively weighted features mapped on to the model structure of the lowest-Tm sequence.

### S.3 Comprehensive analysis of activation patterns for features

In the main text (Fig. 6), we mapped feature activations onto representative structures. To verify whether these activation patterns are conserved at specific positions (AHo numbering) across the entire dataset and how their intensity correlates with Tm ranking, we performed a heatmap analysis. We visualized the activations for the eight key features discussed in Fig. 6 (including SFT positive/negative and pre-trained features) across all 567 sequences sorted by Tm.

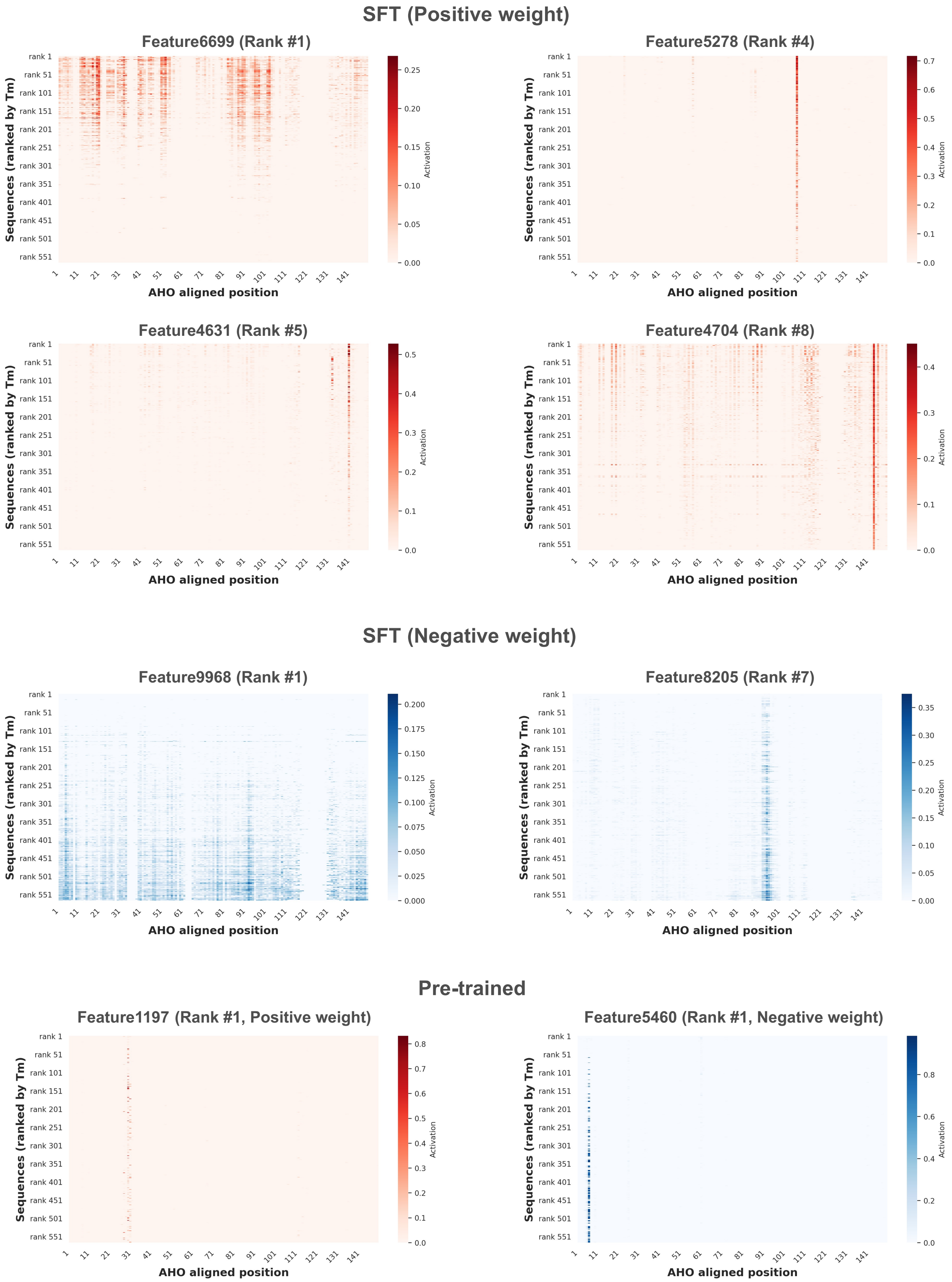


**Figure S5. Heatmaps of feature activation across all AHo-aligned sequences for key features.** Heatmaps showing the activation strength for the eight key features identified in Figure 6 across all 567 NbThermo sequences aligned via AHo numbering. The y-axis represents sequences sorted by Tm in descending order (Rank 1 corresponds to the highest Tm; Rank 567 to the lowest), and the x-axis represents the AHo aligned positions (1-149). Color intensity indicates normalized activation strength.
